# Resolving plasmid-encoded carbapenem resistance dynamics and reservoirs in a hospital setting through nanopore sequencing

**DOI:** 10.1101/2025.08.20.671332

**Authors:** Ela Sauerborn, Rhys T. White, Anna-Lena Kalteis, Daniel Gygax, Ebenezer Foster-Nyarko, Nina Wantia, Friedemann Gebhardt, Lara Urban

## Abstract

The growing resistance of Enterobacterales to last-resort antibiotics such as carbapenems puts a significant burden on healthcare systems, also due to plasmids driving a rapid spread of carbapenem resistance. We here evaluate the use of long-read nanopore sequencing to investigate carbapenem resistance dynamics and the role of plasmid transfers and environmental reservoirs in the hospital setting. Over 13 months, routine clinical diagnostics identified recurring isolates of carbapenem-resistant *Citrobacter* species carrying *Klebsiella pneumoniae* carbapenemases (KPC) and/or OXA-48-like carbapenemases from patient screening and hospital drain samples. While routine diagnostic approaches provided limited insights into the carbapenem resistance dynamics, we show that near-complete *de novo* assembly of chromosomes and plasmids by long-read nanopore sequencing allowed for high-resolution strain identification, plasmid profiling, and antibiotic resistance gene detection. Notably, genomically nearly indistinguishable *Citrobacter freundii* of the high-risk sequence type ST91 genomes were recovered from screening samples collected in the same hospital room one year apart. We further provide evidence of a KPC-2 encoding IncN plasmid that is likely to have spread across bacterial species and between patient and drain isolates, which emphasizes the role of contaminated drains in the persistence and dissemination of resistances within the hospital environment. Overall, this study demonstrates the value of long-read nanopore sequencing for uncovering the complex dynamics of carbapenem resistance spread and persistence in the hospital setting, and its potential implications for Infection Prevention and Control.

**Impact statement:** This study demonstrates how long-read nanopore sequencing can resolve the complex dynamics of plasmid-mediated antimicrobial resistance in clinical and environmental samples within the hospital setting. By linking patient- and drain-derived isolates through near-complete *de novo* assemblies, we reveal hidden reservoirs and dynamics behind the persistence of cabapenem resistance over extended time periods. This work shows how long-read sequencing approaches can uncover resistance dynamics that are missed using standard diagnostic methods, with implications for infection control and surveillance.

**Data Summary:** The study sequences are available at the National Center for Biotechnology Information (NCBI) under BioProject accession number PRJNA1297122. The raw sequence read data is available at NCBI sequence read archive (SRA (https://www.ncbi.nlm.nih.gov/sra)) under accession numbers SRR34727947-59. The chromosomal assemblies of all ST91 strains are available at NCBI GenBank under the Biosample accession numbers SAMN50449475-80. All other supporting data are provided in the article and supplementary data files.

## Introduction

Antimicrobial resistance, particularly to last-resort antibiotics such as carbapenems, is a growing health concern (1,2). Infections with carbapenem-resistant Enterobacterales (CRE), including *Citrobacter* spp., present significant clinical challenges due to limited therapeutic options leading to delayed or ineffective treatment and increased morbidity and mortality (1). The dissemination of carbapenem resistance is primarily driven by horizontal gene transfer via plasmids (3–6). This mechanism enables the rapid spread of resistance across bacterial species boundaries, complicating efforts to control the plasmid-mediated resistance spread in healthcare settings (5–7). Understanding plasmid-encoded resistance dynamics is therefore crucial for developing effective containment strategies.

Environmental reservoirs of CREs within healthcare facilities serve as persistent sources of CRE transmission and amplification. Hospital sink and shower drains have been identified as particularly problematic reservoirs, providing optimal conditions for CRE survival and proliferation (8–10). In clinical settings, the impact of a carbapenem resistance spread is largely determined by how quickly it is detected, as this directly affects the speed and effectiveness of the IPC response (11). Rapid and accurate genomic surveillance and plasmid characterization of both patient and environmental samples are therefore essential for understanding spread and persistence mechanisms of carbapenem resistance (6,7,12–14). However, many clinical microbiology laboratories lack in-house genomic capabilities and instead rely on phenotypic testing methods that cannot resolve resistance mechanisms at the genetic level, or on external sequencing services that provide results too slowly to inform rapid IPC responses. *In situ* nanopore sequencing offers significant advantages for real-time genomic surveillance in the clinical setting. Its capacity for long-read sequencing enables the resolution of complex genomic structures, including complete plasmid sequences, while its rapid turnaround time supports timely IPC decision-making. These capabilities make nanopore sequencing particularly valuable for disentangling complex epidemiological relationships in healthcare-associated outbreaks (15–19).

In this study, we conducted a retrospective genomic investigation of a carbapenem-resistant *Citrobacter* spp. cluster in an internal medicine ward using whole-genome nanopore sequencing. Over a 13 month period, routine laboratory diagnostics detected *Klebsiella pneumoniae* carbapenemases (KPC)-producing and/or OXA-48-like carbapenamase producing *Citrobacter* spp. in patient screening and environmental samples from sink and shower drains. We here show that conventional diagnostic approaches based on phenotypic testing and lateral flow assays could not sufficiently resolve the dynamics behind this cluster of carbapenem-resistant *Citrobacter* spp. In contrast, long-read nanopore sequencing, allowed us to characterize complex carbapenem resistance dynamics and identify critical environmental reservoirs contributing to persistence and transmission within the hospital setting.

## Methods

### Sample collection

From February 2024 to March 2025, recurring carbapenem-resistant *Citrobacter* spp. carrying KPC and/or OXA-48-like carbapenemases, were repeatedly detected in patient screening and environmental samples from an internal medicine ward at the Technical University of Munich (TUM) University Hospital in Munich, Germany. Relevant isolates were detected in ten screening samples from patients (rectal swabs and one urine sample) and in three environmental (sink and shower drains) samples. Rectal swab collection for CRE detection is routinely conducted as part of the admission screening at the hospital. Briefly, rectal swabs (Becton Dickinson GmbH, Heidelberg, Germany) were collected by insertion to a depth of 2– 3 cm with threefold rotation. One mid-stream urine sample was included in this study due to the phenotypic detection of carbapenem-resistant *Citrobacter* spp. (see below for details on phenotypic tests). All patient samples were obtained from individual patients. The samples were plated out on BD® MacConkey (Becton Dickinson GmbH, Heidelberg, Germany) and Thermo Scientific™ Brilliance™ ESBL agar plates and incubated at 37°C for 20-24hours. Environmental drain sample collection was conducted in October 2024 for all rooms in question. At the time of sampling, the sanitary facilities were cleaned as part of the daily routine in accordance with the hospital’s internal cleaning and disinfection protocol by using a limescale-removing sanitary cleaner (Milizid®, DR.SCHNELL GmbH & Co. KGaA, Munich, Germany). No specific treatment of the drains was carried out prior to drain sampling. For sample collection, a 50 mL bladder syringe was fitted with a suction catheter and inserted into each drain. The system was repeatedly aspirated and flushed to mix the contents and 20 mL of this mixture was drawn from the drain and transferred it into a tryptic soy broth (TSB) enriched with disinhibitor to deactivate residual disinfectants (Thermo Scientific™) for incubation. After 20-24 hours, 100 µL of the broth were plated out on BD® Columbia Blood, MacConkey (Beck Dickinson GmbH, Heidelberg, Germany) and Thermo Scientific™ Brilliance™ ESBL agar plates; If no growth was detected in the broth after 48 hours, the plating was repeated. Environmental samples showing no growth in the broths or on the plates thereafter were declared negative.

### Established diagnostics

For established laboratory diagnostics, species identification was conducted from a single colony forming unit (CFU) of bacterial isolates with growth on ESBL agar plates using Matrix-assisted laser desorption ionization time-of-flight mass spectrometry instructions (MALDI-TOF MS, Bruker Daltronics GmbH, Leipzig, Germany), as per the manufacturer’s instructions. Phenotypic antibiotic susceptibility testing was performed using VITEK 2 (BioMérieux, Marcy l’Etoile, France) on pure subcultures. For this, up to three bacterial CFUs were transferred to a saline tube to generate a homogenous suspension with a density equivalent to 0.5 McFarland. Subsequently, minimum inhibitory concentrations (MICs in mg/L) of the bacterial isolates were determined using the VITEK 2 gram-negative (AST-GN69) card, and the carbapenem susceptibility results interpreted according to the European Committee on Antimicrobial Susceptibility Testing (EUCAST) guidelines (20). Carbapenemases were identified using a multiplex immunochromatography assay consisting of lateral flow assays (O.K.N.V.I Resist-5) that detects the presence of KPC-2, OXA-48-like, NDM, VIM, IMP-carbapenemases, according to manufacturer’s instructions. After completing these routine diagnostic steps, five to ten CFUs of each isolate were stored at −80 °C in glycerol stocks for future use.

### DNA extraction and nanopore sequencing

Stored isolates were grown overnight at 37 °C on BD® Columbia blood agar plates. Per isolate, DNA was extracted in a spin-column-based DNA purification approach. For this, 10-20 CFUs of overnight bacterial cultures were resuspended in 1mL phosphate-buffered saline (PBS). A 100μL aliquot of this suspension was transferred to a fresh 1.5mL Eppendorf tube and adjusted to 1mL with PBS, mixed gently, and centrifuged at 12,000 x g for 2 min. The supernatant was discarded, and genomic DNA was extracted from the resulting pellet with the Qiagen DNeasy Blood & Tissue kit (Qiagen, Hilden, Germany), following the manufacturer’s Gram-negative bacteria protocol (21). We added 40μL of proteinase K and extended the lysis incubation to 2 hours to increase yield and purity of the DNA extracts.

The resulting DNA extracts were quantified using a Qubit 4 Fluorometer based on DNA-specific fluorescent dyes (dsDNA HS kit), and to NanoDrop One Spectrophotometer measurements of total nucleic acids based on UV-absorbance at 260nm. In a subset of DNA extracts, we observed a discrepancy between Qubit and NanoDrop measurements (Table S1) indicated excess residual RNA (Table S1) , which can impact library preparation and contribute to background sequencing signal. Indeed, high levels of failed sequencing reads (Fig. S1a) and unavailable nanopores (Fig. S1c) when sequencing these extracts were detected preventing any further robust downstream analysis. We therefore treated these DNA extracts with high RNA content post hoc by adding 4µl of DNase free RNase (10mg/ml, Thermo Fisher Scientific, Watham, USA) directly to 50 µl of DNA extracts, followed by vortex mixing and a 15-minute incubation step at room temperature. As DNA yield was reduced after RNAse treatment (Table S1), the resulting DNA extracts were further purified and concentrated with the Zymo DNA Clean & Concentrator-5 kit (Zymo Research, Irvine, USA) according to the manufacturer’s instructions. The post hoc RNase treatment substantially removed residual RNA and resulted in better sequencing outcomes (Table S1; Figs. 1b & 1d). However, as it also reduced DNA yield and purity (Table S1), we only recommend its post hoc application on non-precious DNA extracts. Based on these observations, an RNase step was incorporated into the protocol of subsequent DNA extractions after the proteinase K incubation step, following the manufacturer’s instructions (Qiagen, Hilden, Germany).

**Fig. 1.**
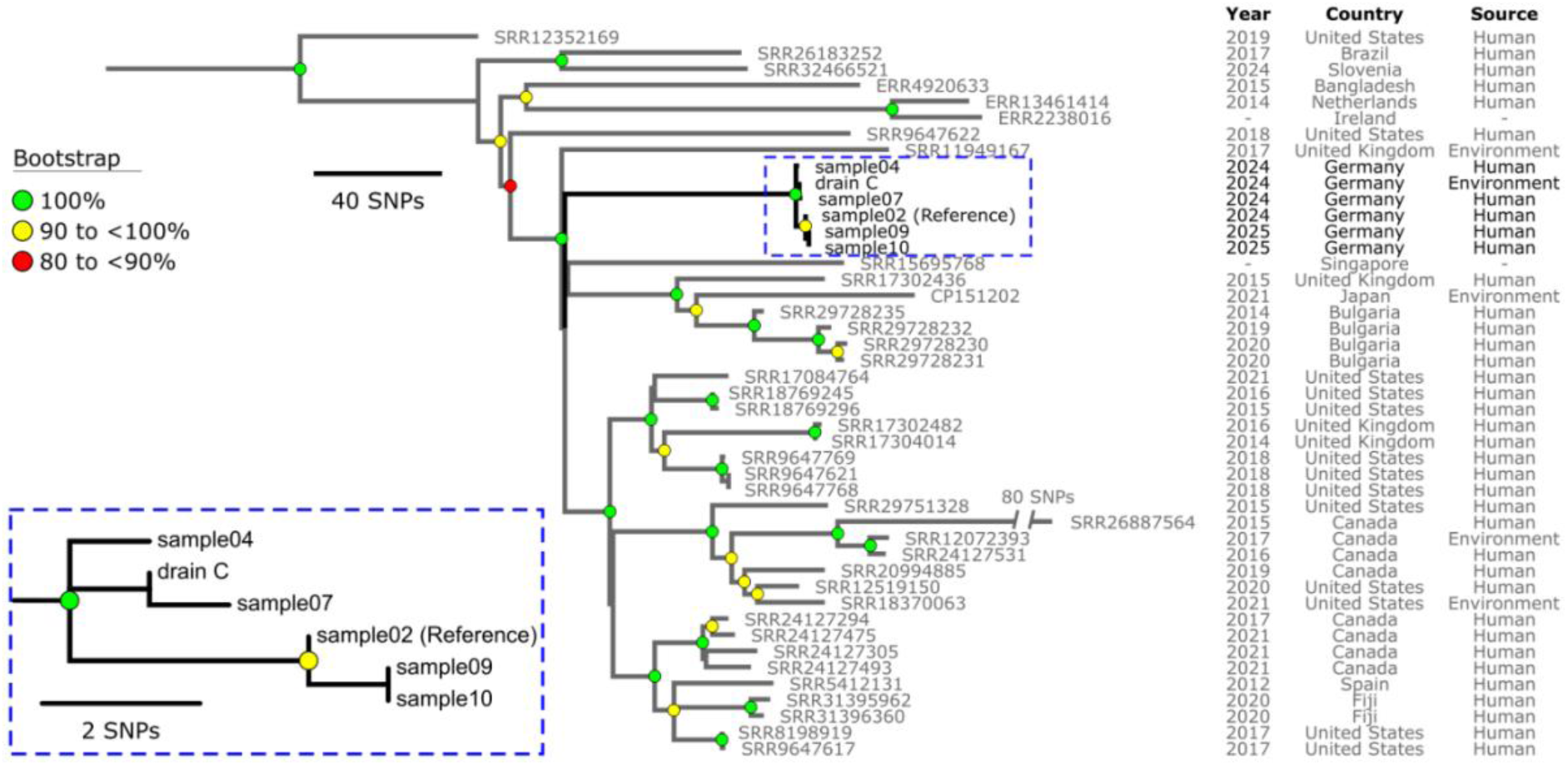
Maximum-parsimony phylogeny of the global *Citrobacter freundii* sequence type ST91 lineage. The phylogeny was inferred from 1,772 non-recombinant orthologous biallelic core-genome single-nucleotide polymorphisms (SNPs) from 45 genomes. SNPs were derived from a core-genome alignment of ∼4,124,900 bp and were called against the chromosome of sample02 (GenBank accession number SAMN50449475 The consistency index for the tree was 1.0. SNP density filtering in SPANDx (excluded regions with three or more SNPs in a 10 bp window). The phylogenetic tree was rooted according to the ST62 outgroup, which has been omitted for visualization (Methods).

Nanopore sequencing libraries of all DNA extracts were generated using the SQK-RBK114 Rapid Barcoding Kit and sequenced on R10.4.1 flow cells for 16-20 hours using an Oxford Nanopore Technologies MinION MK1d device. We used two barcodes per DNA extract to account for imbalanced sequencing outputs across barcodes. 90ng to 200ng DNA in 10µl of DNA extract per barcode were used as input for nanopore library preparation (as recommended by the manufacturer), depending on the input DNA concentration.

### Nanopore data basecalling and quality control

Basecalling, genome assembly, AMR gene detection and plasmid typing were conducted on a portable laptop with an 8 GB NVIDIA GeForce RTX 4070 GPU, 16 GB 5200 MHz RAM, and an Intel i7-13800H CPU with 14 cores and 20 threads. The corresponding data analyses protocols are provided at https://github.com/ElaSrbrn/Carbepenem-resistant-Enterobacterales-Plasmid-clustering-pipeline. The raw nanopore data was basecalled using Dorado v5.0 and the SUP basecalling model (<u>dna_r10.4.1_e8.2_400bps_sup@v5.0.0</u>). We used Porechop v0.2.3 (https://github.com/rrwick/Porechop, accessed 28 June 2025) to trim the adapter sequences and filtered out low-quality reads (Q < 9) and short sequences (< 200 bases) using Nanofilt v2.8.0 (https://github.com/wdecoster/nanofilt, accessed 28 June 2025)(22). Sequencing summaries were generated using Seqkit v2.10.0 (https://github.com/shenwei356/seqkit, accessed 28 June 2025) (23).

### Nanopore-based *de novo* assembly and genotyping

We generated *de novo* assemblies using Flye v2.9.5 (24) for high quality long reads (nano-hq) followed by polishing with Medaka v2.01 (https://github.com/nanoporetech/medaka, accessed on 22 July 2025), in the bacterial mode (--bacteria). We assessed assembly coverage using SAMtools depth v1.18 (25), ensuring that all assemblies had a minimum median chromosomal coverage of 40× (26). Contig circularity was visually assessed using bandage v0.9.0 (27). The chromosomes were reoriented to start at the *dna*A gene using dnaapler v1.2.0 (28).

We then analysed our polished *de novo* assemblies using the Pathogenwatch v2.3.1 platform (accessed 12 July 2025) for species identification, multi-locus sequence typing (MLST), and resistance gene detection (29). Any discordance between Pathogenwatch and MALDI-TOF MS species calls was investigated with Kraken2 v2.1.4 (30) using default parameters and the National Center for Biotechnology Information (NCBI) nucleotide database (https://www.ncbi.nlm.nih.gov/nucleotide, accessed on 14 January 2025). Additionally, we confirmed the discordance in species identification with the *Citrobacter* spp. database hosted on PubMLST (accessed 12 July 2025) (31).

### Global phylogeny of publicly available *Citrobacter freundii* genomes

We next conducted phylogenetic analyses on publicly available *C. freundii* genomes to identify the species’ global diversity as a basis for our local clustering analyses. For this, Complete *C. freundii* genomes were retrieved from the NCBI nucleotide database (https://www.ncbi.nlm.nih.gov/nucleotide, accessed 24 July 2025). Additionally, paired-end Illumina sequence data for *C. freundii* was obtained from the NCBI sequence read archive (SRA (https://www.ncbi.nlm.nih.gov/sra, accessed 24 July 2025) for sequence data belonging to *C. freundii* using the ‘fasterq-dump’ tool within the SRA Toolkit v3.0.1-ubuntu64 (https://github.com/ncbi/sra-tools, accessed 28 July 2025), restricting searches to whole-genome sequencing, whole-genome amplification, whole-chromosome sequencing, clone-based, finishing, or validation strategies. Raw Illumina sequence reads were *de novo* assembled using Shovill v1.1.0 (https://github.com/tseemann/shovill, accessed on 28 July 2025) with parameters set to estimate the genome size at 5 Mb, remove contiguous sequences (contigs) with a sequence coverage below 20-fold and enable single-cell mode. Assembly metrics were assessed using QUAST v5.0.2 (39) , with genomes removed if assemblies < 4.3 Mb or > 6.2 Mb, had N50 values < 10,000 bp, or sequencing depth < 20×.

The resulting assemblies were input into a Genome Taxonomy Database Toolkit (GTDB-Tk) genome-based taxonomy (GTDB-Tk v2.1.1 with GTDB package R207_v2 (40)). Sequences within the genus *Citrobacter* and *Salmonella* (g_, genus) were extracted to construct a concatenated reference alignment of 120 bacterial marker genes. The taxonomic tree was inferred using maximum-likelihood approximation with FastTree v2.1.7 (41) under the WAG model (42) of protein evolution with gamma-distributed rate heterogeneity (43) (+ GAMMA). Genomes clustering outside of *C. freundii* species lineage were removed from downstream analyses. *In silico* MLST was done using MLST v2.23.0 (https://github.com/tseemann/mlst, accessed on 28 July 2025) with default settings to query the assemblies against the *Citrobacter* spp. typing database (44) hosted on PubMLST (45) (local database updated 19 March 2024). A total of 2,028 *C. freundii* genome assemblies (quality filtered (*n*= 1,861); complete (*n*= 167)) were aligned to create a core-genome alignment using Parsnp v1.7.4 (46), with the reference being the chromosome of CFTMDU (GenBank: CP151202), to identify single-nucleotide polymorphisms (SNPs). Resulting SNP alignments were used to reconstruct phylogenies. We used RaxML v8.2.12 (47) to build phylogenetic trees using the maximum-likelihood method with GTR-GAMMA correction (optimizing 10 distinct, randomized maximum-parsimony trees). The phylogenetic trees were visualized using FigTree v1.4.4 (http://tree.bio.ed.ac.uk/software/figtree/, accessed 28 July 2025).

### Local genomic clustering of *Citrobacter freundii* sequence type (ST)91

We used the global *C. freundii* phylogeny to identify globally available C*. freundii* sequence type ST91 genomes, their closest phylogenetic outgroup, and investigate global delineation of the ST91 lineage. Genomic variants between all publicly available and our local ST91 genome were identified using the SPANDx v4.0.4 pipeline (48), mapping reads to lineage-specific reference chromosomes. SNPs within regions of high-density clusters (≥3 SNPs in 10 bps window), mobile genetic elements, and predicted recombination sites (identified using Gubbins v3.3.5 (49)) were excluded. Sites were excluded if an SNP was called in regions with less than half or greater than 3-fold the average genome coverage on a genome-by-genome basis. This analysis defines a core genome as regions estimated to the nearest 100 bp with ≥95 % coverage in all genomes, as calculated using the BEDTools v2.28.0 (50) coverageBed module within the SPANDx pipeline. The pairwise SNP distances were determined using snp-dist v0.6.3 (https://github.com/tseemann/snp-dists, accessed 22 July 2025). Maximum-parsimony trees were reconstructed from the orthologous biallelic core-genome SNP alignments using the heuristic search feature of PAUP v4.0a (51).

### Carbapenemase gene detection and plasmid characterisation

We identified contig-specific carbapenem resistance genes in our generated nanopore assemblies using AMRFinderPlus v4.0.3 (https://github.com/ncbi/amr, accessed 28 July 2025) with database version 2024-10-22.1 (52,53). Carbapenemase-encoding contigs were functionally annotated with MOB-suite v3.1.8 (54). The MOB-typer module (54) was then applied to the contigs identified as plasmids to detect key mobility determinants to estimate the potential for horizontal gene transfer. We visualised the plasmid annotation using the mobileOG-db (55) integrated in ProkSee (56) with the carbapenemase positions confirmed using the CARD database (57).

### Plasmid similarity analyses

Plasmid similarity analyses were conducted for carbapenemase-encoding plasmids with potential for interspecies transfer. The plasmid clustering followed a stepwise approach starting with pairwise MinHash (Mash) based approximation of Jaccard distances for shared k-mers using MASH v1.4.5 (58). We applied a Mash distance threshold of 0.001 to define highly similar plasmid sequences (4,59), and excluded those plasmids that did not fulfil this threshold criteria with any other plasmid from further analyses. We next assessed the structural relatedness of the remaining plasmids by applying the Double-Cut-and-Join (DCJ)-indel model using pling v1.0.1 with a containment threshold of 0.3 (recommended for recent transmission events) and default threshold of 4 DCJ and indel operations needed to transform one plasmid into another (59). Additional exploratory linear comparative analyses of the assembled plasmids were performed using the ProkSee BLASTn module (56).

## Results

### Species identification and antimicrobial resistance detection with established diagnostics

Established laboratory diagnostics classified isolates as *Citrobacter farmerii* (*n*=2) and *C. freundii complex* (*n*=11) using MALDI-TOF MS and identified carbapenem resistance in all isolates using VITEK 2 (Methods; Table 1). Subsequent lateral flow assays identified six KPC-positive isolates, six isolates positive for both OXA-48-like- and KPC carbepenemases, and one isolate positive for OXA-48-like- and VIM carbapenemases (Methods; Table 1). Notably, room D was the only room with patient occupancy that yielded CRE-negative drain samples during environmental sampling. Follow-up drain sampling in March 2025 after identification of the CRE positive samples 9 and 10 from room D again returned negative results for CRE growth (Table 1).

**Table 1.** Bacterial species and antimicrobial resistance mechanism identifications across all isolates from patient (1–10) and drain (A-D) samples using established diagnostics and nanopore sequencing-based de novo assemblies. For each sample, sampling source, date, and room are indicated. For nanopore sequencing, sequence types (ST) according to MLST and resistance gene location (chromosome- or plasmid-encoded with replicon types IncN or IncL/M) are additionally indicated in brackets.

| Sample |  |  | Bacterial species identification |  | Antimicrobial resistance mechanism identification |  |
| --- | --- | --- | --- | --- | --- | --- |
| No | Date | Room | Established diagnostics | Nanopore sequencing | Established diagnostics | Nanopore sequencing |
| 1 | 02/2024 | A | *C. freundii* complex | *Enterobacter mori* | KPC | KPC-2 (IncN) |
| 2 | 03/2024 | B & D | *C. freundii* complex | *C. freundii* (ST 91) | KPC<br>OXA-48-like | KPC-2 (IncN),<br>OXA-48 (IncL/M) |
| 3 | 05/2024 | C | *C. farmerii* | *C. farmerii* (ST 857) | KPC | KPC-2 (IncN) |
| 4 | 06/2024 | A & D | *C. freundii* complex | *C. freundii* (ST 91) | KPC<br>OXA-48-like | KPC-2 (IncN),<br>OXA-48 (IncL/M) |
| 5 | 08/2024 | B | *C. farmerii* | *C. farmerii* (ST 857) | KPC | KPC-2 (IncN) |
| 6 | 09/2024 | C | *C. freundii* complex | *C. portucalensis* | OXA-48-like<br>VIM | OXA-48, VIM (chromosome) |
| 7 | 09/2024 | C | *C. freundii* complex | *C. freundii* (ST 91) | KPC<br>OXA-48-like | KPC-2 (IncN),<br>OXA-48 (IncL/M) |
| 8 | 10/2024 | A & B | *C. freundii* complex | *C. youngae* | KPC | KPC-2 (IncN) |
| 9 | 03/2025 | D | *C. freundii* complex | *C. freundii* (ST 91) | KPC<br>OXA-48-like | KPC-2 (IncN),<br>OXA-48 (IncL/M) |
| 10 | 03/2025 | D | *C. freundii* complex | *C. freundii* (ST 91) | KPC<br>OXA-48-like | KPC-2 (IncN),<br>Oxa-48 (IncL/M) |
| Drain A | 10/2024 | A | *C. freundii* complex | *C. portucalensis* | KPC | KPC-2 (IncN) |
| Drain B | 10/2024 | B | *C. freundii* complex | *C. freundii* (ST 18) | KPC | KPC-2 (IncN) |
| Drain C | 10/2024 | C | *C. freundii* complex | *C. freundii* (ST 91) | KPC<br>OXA-48-like | KPC-2 (IncN),<br>OXA-48 (IncL/M) |
| Drain D | 10/2024 | D | No growth | NA | NA | NA |

### Nanopore sequencing and assembly metrics

Nanopore sequencing resulted in a mean N50 of >7 kb across all filtered sequencing reads (Methods), and an overall high mean quality score ranging from 20.6 to 22.2 across all isolates (Table S2). All *de novo* assemblies of bacterial chromosomes and IncN plasmids exceeded the minimum median coverage of 40× (Table S3). Chromosomal assemblies ranged from 4.8 Mb to 5.3 Mb in length, while IncN plasmid assemblies ranged from 72 kb to 88 kb. Notably, plasmid contigs from the isolates detected in samples 4 and 5 were approximately 10 kb longer than other IncN plasmid assemblies (Table S3).

### Bacterial species identification through genomic data analysis

Comparison of species identification between routine diagnostics and nanopore sequencing revealed important discrepancies and provided enhanced taxonomic resolution (Methods; Table 1). Nanopore sequencing confirmed the identification of two *C. farmerii* isolates initially classified by established diagnostics. However, the isolate from sample 1, initially identified as *C. freundii* complex, was reclassified as *Enterobacter mori* by all computational annotation tools, including Pathogenwatch, Kraken 2, and PubMLST (Methods; Table 1). For the remaining isolates identified within the *C. freundii complex*, nanopore sequencing provided species-level resolution that established diagnostics could not achieve: The isolates from sample 7 and drain sample A were identified as *Citrobacter portucalensis*, sample 9 was resolved to *Citrobacter youngae*, and the remaining seven isolates to *C. freundii*, which are all part of the *C. freundii complex* (Methods; Table 1). Multi-locus sequence typing (MLST) revealed six of the seven *C. freundii* isolates belonged to the same high-risk *C. freundii* strain ST91 (60) (Methods; Table 1), indicating clonal relatedness.

### Global diversity and local clustering of *Citrobacter freundii* ST91

To contextualise our ST91 isolates within global diversity, we explored the relatedness of the six *C. freundii* ST91 isolates through global phylogenetic and subsequent local variant calling analyses. To correctly root the ST91 phylogeny, we first examined the overall population structure of *C. freundii* (Fig. S2) by including 2,028 publicly available *C. freundii* assemblies identified and filtered through our assembly-based taxonomic classification (Methods, Fig. S2, Fig. S3). Maximum-likelihood phylogenetic analysis identified 39 ST91 genomes from 13 countries spanning 2012 to 2024, comprising 33 human-derived isolates and four environmental samples from sources such as sewage, hospital drains, river and wastewater (Supplementary data 1). Clinical samples types showed diverse anatomical origins, with urine (*n* = 11) being most common, followed by wound/abscess samples (*n* = 6). ST62 served as the closest outgroup to ST91, confirming that ST91 represents a well-defined lineage within the *C. freundii* species complex.

Integration of the six *C. freundii* ST91 bacterial chromosomes sequenced in this study to the 39 publicly available ST91 genomes revealed striking epidemiological patterns (Fig. 1). The global ST91 phylogeny is genomically diverse, with pairwise SNP distances ranging from 0 to 396 SNPs (median = 153; interquartile range (IQR) = 116 to 189) (Supplementary data 1). In contrast, the six isolates from this study (2024 to 2025) form a highly clonal monophyletic cluster, with pairwise SNP distances ranging from 0 to 6 SNPs (median = 4; IQR = 1.5 to 5 SNPs; Table 2). Within this local cluster, pairwise SNP distances ranged from 0 to 6 SNPs (median = 4; IQR = 1.5 to 5 SNPs; Table 2), indicating recent transmission or ongoing circulation within the hospital setting.

**Table 2.** Pairwise SNP-based distance matrix between all TUM University Hospital *Citrobacter freundii* ST91 bacterial chromosomes (Methods).

| Sample | 2 | 4 | 7 | Drain C | 9 | 10 |
| --- | --- | --- | --- | --- | --- | --- |
| 2 | - | 4 | 5 | 4 | 1 | 1 |
| 4 | 4 | - | 3 | 3 | 5 | 5 |
| 7 | 5 | 3 | - | 1 | 6 | 6 |
| Drain C | 4 | 5 | 1 | - | 5 | 5 |
| 9 | 1 | 5 | 6 | 5 | - | 0 |
| 10 | 1 | 5 | 6 | 5 | 0 | - |

Within our cluster, isolates from samples 9 and 10 are genomically indistinguishable across the core genome, and the smallest pairwise SNP distance (1 SNP) was observed among the ST91 strains from samples 2, 9, and 10. Thus, the first ST91 index isolate of this local cluster (sample 2), forms a tight phylogenetic sub-branch with ST91 isolates from sample 9 and 10, reflecting their close relatedness (Fig. 1). The ST91 from sample 7 also differed by a single SNP from drain C further supporting the hypothesis of environmental reservoir-mediated transmission. All observed SNP distances (Table 2) fell well within established thresholds for defining epidemiologically linked clusters in Enterobacterales (60,61).

### Comprehensive antimicrobial resistance characterization through genomic data analysis

Nanopore sequencing confirmed all carbapenemases detected by routine diagnostics while providing enhanced genetic context and subtype identification (Methods; Table 1). All carbapenemases detected by routine diagnostics were confirmed with the nanopore-based workflow. The *de novo* assemblies additionally identified the subtype and genetic context of each carbapenemas. All KPC-positive isolates across four different bacterial taxa carried the same KPC subtype KPC-2, which were all located on a replicon type IncN plasmid. These IncN plasmids belonged to the same MOB Cluster AA552, and shared conserved mobilisation features such as a MOBF relaxase, MPF_T mating-pair formation system, and a MOBF origin of transfer, confirming their classification as conjugative plasmids by MOB-suite (54).

OXA-48-like-carbapenemases were confirmed as OXA-48 subtypes in all positive isolates. Notably, genetic context varied by bacterial species: OXA-48 was chromosomally encoded in *C. portucalensis* (sample 6), while all *C. freundii* ST 91 isolates carried OXA-48 on conjugative IncL/M plasmids (Table 1).

### Plasmid clustering reveals complex transmission networks

Given the multi-species distribution of KPC-2-carrying IncN plasmids across different environmental and clinical sources, we focused plasmid clustering analyses on the IncN plasmids to better understand their dynamics (Method,Table 1). Mash distance analysis revealed distinct clustering patterns, with plasmids from sample 3 (*C. farmerii*) and drain sample B (*C. freundii* ST 18) showing a pairwise distance of >0.001 with all other IncN plasmids and were thus excluded from further plasmid clustering analyses (Table 3). The IncN plasmid from sample 1 (*E. mori*) clustered most closely with plasmids from sample 8 (*C. youngae*) and drain sample A (*C. portucalensis*), all associated with room A and sampled 8 months apart (Table 1; Table 3; Fig. S4b). This spatial-temporal clustering suggests room-specific plasmid circulation. Similarly, five of six *C. freundii* ST91 strains (found in samples 2, 7, 9, 10, and drain sample C) formed a tight plasmid cluster with low Mash distances (Table 3). The IncN plasmid from *C. freundii* ST91 from sample 4 represented a notable exception, differing in length by approximately 10 kb from other ST91 IncN plasmids (Table S3). Comparative genomic analysis revealed an additional Tn3 transposon insertion partially accounting for this length difference (Fig. S4c). This variant plasmid clustered most closely with the IncN plasmid from sample 5 (*C. farmerii*), with which it shared similar length characteristics (Table 3, S3).

**Table 3.** Pairwise Mash distances of all IncN plasmids detected in bacterial isolates across samples. Pairwise distances at a clustering threshold of >0.001 are marked in grey (Methods).

| Sample & Room | 2<br>B/D | 3<br>C | 4<br>A/D | 5<br>D | 7<br>C | 8<br>A/B | 9<br>D | 10<br>D | Drain<br>A | Drain<br>B | Drain<br>C |
| --- | --- | --- | --- | --- | --- | --- | --- | --- | --- | --- | --- |
| 1 A | 0.0004 | 0.0024 | 0.0025 | 0.0025 | 0.0004 | 4.77e-05 | 0.0004 | 0.0004 | 7.16e-05 | 0.010 | 0.0004 |
| 2 B/D | x | 0.0029 | .0029 | 0.0028 | 0 | 0.0004 | 0 | 0 | 0.0004 | 0.009 | 2.39e-05 |
| 3 C | 0.0024 | x | 0.005 | 0.005 | 0.0029 | 0.0024 | 0.0028 | 0.0028 | 0.0024 | 0.013 | 0.0028 |
| 4 A/D | .0029 | 0.005 | x | 9.55e-05 | 0.0029 | 0.0025 | 0.0028 | 0.0028 | 0.0025 | 0.012 | 0.0029 |
| 5 D | 0.0028 | 0.005 | 9.55e-05 | x | 0.0029 | 0.0025 | 0.0029 | 0.0029 | 0.0025 | 0.012 | 0.0029 |
| 7 C | 0 | 0.0029 | 0.0029 | 0.0029 | x | 0.0004 | 0 | 0 | 0.0004 | 0.009 | 2.38 e-05 |
| 8 A/B | 0.0004 | 0.0024 | 0.0025 | 0.0025 | 0.0004 | x | 0.0004 | 0.0004 | 7.16e-05 | 0.010 | 0.0004 |
| 9 D | 0 | 0.0028 | 0.0028 | 0.0029 | 0 | 0.0004 | x | 0 | 0.0004 | 0.009 | 2.38e-05 |
| 10 D | 0 | 0.0028 | 0.0028 | 0.0029 | 0 | 0.0004 | 0 | x | 0.0004 | 0.009 | 2.38e-05 |
| Drain A | 0.0004 | 0.0024 | 0.0025 | 0.0025 | 0.0004 | 7.16e-05 | 0.0004 | 0.0004 | x | 0.010 | 0.0005 |
| Drain B | 0.009 | 0.013 | 0.012 | 0.012 | 0.009 | 0.010 | 0.009 | 0.009 | 0.010 | x | 0.009 |
| Drain C | 2.39e-05 | 0.0028 | 0.0029 | 0.0029 | 2.38 e-05 | 0.0004 | 2.38e-05 | 2.38e-05 | 0.0005 | 0.009 | x |

All plasmids clustering by Mash distance (<0.001) also satisfied DCJ-indel clustering criteria (DCJ-indel distance ≤4; Fig. 2, Table S4 ), confirming that sequence similarity correlated with structural relatedness. However, some discordances between DCJ and Mash distances highlighted the complexity of plasmid similarity analyses—for instance, IncN plasmids from isolates 1 and 4 showed zero DCJ distance despite moderate Mash distance, indicating structural conservation despite sequence divergence (Table S4, Fig. S4c).

**Fig. 2.**
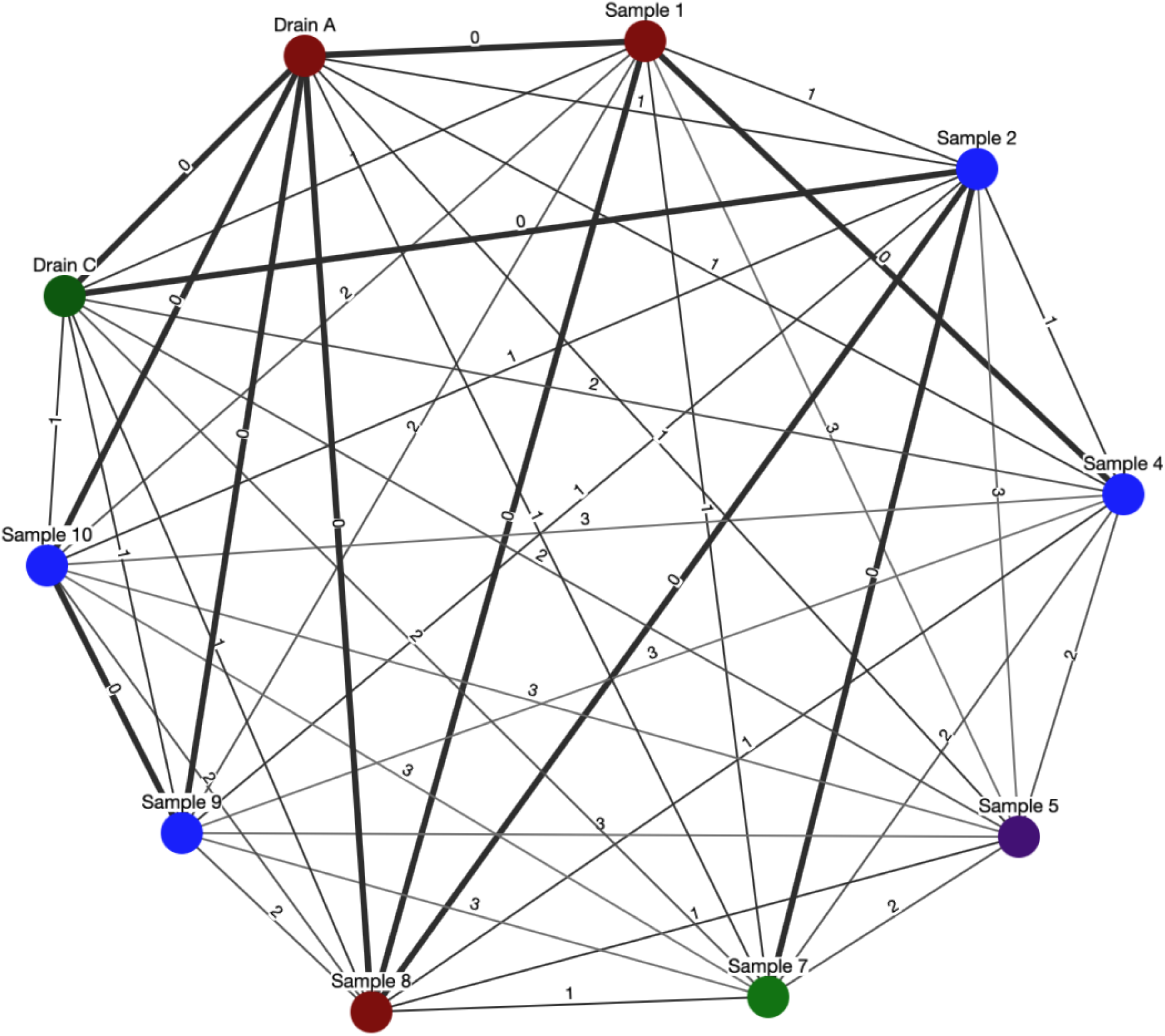
DCJ-indel clustering network of IncN plasmids with nodes representing individual samples colored according to their room location (red: room A, violet: room B, green: room C, blue: room D, see Supplementary Table 4). Edges between nodes indicate the DCJ-indel distance, with bold edges representing a distance of zero. IncN plasmids from samples 3 and drain sample B have been excluded due to a Mash distance above 0.001.

## Discussion

We highlight the relevance of long-read whole-genome sequencing for uncovering bacterial strain- and plasmid-mediated carbapenem resistance dynamics between patient and environmental reservoirs in the hospital, which would otherwise be missed by established diagnostics. In our example, routine culture and susceptibility testing flagged a rise in carbapenem-resistant *Citrobacter* spp. on an internal medicine ward within a timeframe of 13 months. However, the information provided by these established diagnostic tools could not discriminate whether this trend reflected i) a clonal spread of a single *Citrobacter* strain, ii) a horizontal transfer of distinct plasmids, iii) a gain or loss of OXA-48 within a shared KPC-encoding plasmid, or iv) a random accumulation of unrelated carbapenem-resistant *Citrobacter* spp. In contrast to established diagnostic procedure, whole-genome sequencing by long-read nanopore technology provided higher-resolution taxonomic profiles, allowing us to exclude unrelated subspecies or strains from further downstream clonal analyses and to accurately group those with greater chromosomal similarity (63). By retrospectively generating *de novo* assemblies from these bacterial isolates using nanopore sequencing, we were able to resolve resistance dynamics at both chromosomal and plasmid levels, and we showed that the accumulation of CREs was a mixture of a circulating *C. freundii* ST91, and of a plasmid-mediated resistance spread across bacterial species and the hospital environment.

More specifically, we identified a *C. freundii* ST91 cluster comprising six isolates from patient screening samples and one from a drain sample. Notably, this cluster was genetically distinct from other global *C. freundii* ST91. Within our hospital cluster itself, we found two identical isolates collected within the same month (sample 9 and 10), differing by only one SNP from the index ST91 isolate (sample 2), which had been sampled 12 months earlier in the same room. As this SNP-based distance is below the expected core-genome point-mutation rate in actively replicating Enterobacterales (62,64), such genomic stasis could be indicative of a low-replication reservoir rather than continuous spread over more than 12 months. One plausible source for this would be environmental biofilms in the hospital setting as found in drains, where CRE embedded in biofilms replicate slowly, accrue few mutations, and can intermittently slough off biofilm fragments that then reseed with nearly identical clones (9,10,65). However, while the other rooms implicated in this study were CRE-positive, even repeated drain cultures from this index room remained negative. This finding might highlight limitations of culture-based surveillance for dormant biofilms, where an extension of the liquid culture enrichment beyond 48 hours, or direct metagenomic sequencing of environmental samples could potentially increase sensitivity. Alternatively, the environmental reservoir might have been located outside of the drain system, which emphasizes the need for more extensive environmental surveillance in the clinical setting. Independently of the scenario, the fact that the index ST91 isolate formed a subcluster with isolates from samples collected 12 months later and is more distantly related to the intermediate ST91 isolates suggests that the true root of the bacterial transmission chain in the internal medicine ward might have been missed. This highlights the common challenges of retrospectively inferring transmission events based on a limited sample size sampled across long time intervals (66).

Beyond chromosomal insights, long-read nanopore sequencing data generated near-complete *de novo* assemblies of plasmids, clarifying whether carbapenem resistance genes were encoded on a plasmid or chromosome, and uncovering the relatedness of plasmids between bacterial species and sampling sites. These *de novo* assemblies identified the role of a KPC-2 carrying IncN plasmid in the described rise of carbapenem resistant isolates. The sequence-level and structural comparisons of the IncN assemblies revealed evidence of interspecies plasmid transfer across four bacterial taxa, and of dissemination between patient and environmental reservoirs over eight months, thereby allowing us to confirm (or rule out) potential epidemiological links. The interpretation of plasmid dynamics, however, requires caution since plasmids show high genomic plasticity and lack a defined molecular clock (such as SNP-based distance for bacterial chromosomes) to infer plasmid relatedness. Thus, any threshold-based inference must still be carefully cross-checked against epidemiological context before conclusions are drawn. To mitigate this uncertainty, we assessed plasmid relatedness with two complementary metrics: sequence similarity (Mash distance) and structural similarity (DCJ-indel distance). This approach captures the high plasticity of plasmids that may undergo substantial structural changes with minimal changes in sequence content and *vice versa* (59). Furthermore, we applied stringent cut-offs as recommended in previous studies (4,59) to minimize the risk of including false positive links between plasmids. Our results highlight that plasmids can readily cross bacterial species and system boundaries in the hospital setting, with implications for clinical surveillance and intervention strategies, which should move beyond single-species analyses and explicitly account for the fluid, modular evolution of plasmids. The likely spread of IncN plasmids between hospital environmental and patient-derived isolates (or *vice versa*) might again point towards the role of biofilms as carbapenemase reservoirs in the clinical setting.

Overall, we provide laboratory and computational guidelines for plasmid-mediated resistance dynamic detection and chromosomal clustering through nanopore whole-genome sequencing in clinical settings. Our findings reveal the limitations of relying solely on routine culture-based diagnostics to capture the complexity of carbapenem resistance dynamics, and underscore the value of nanopore sequencing in resolving both chromosomal and plasmid-level relationships between patient and environmental reservoirs in the clinical setting.

## Author contributions

ES and LU conceptualised the project. ES and LU designed the DNA extraction and sequencing protocols. ES and DG designed the RNA depletion experiment and sequencing comparisons. ES conducted the bioinformatic analysis including data curation, formal analysis, and visualisation, under RTW, LU and EFN supervision. RTW supervised and conducted the chromosomal clustering analysis and EFN consulted on genomic data analysis and interpretation and edited manuscript drafts. FG and NW provided samples and laboratory equipment for phenotypic testing. AK and FG supervised and synthesized the clinical and drain sample collection and metadata overview, as well as the established laboratory diagnostics. ES and LU wrote the manuscript with input from all co-authors.

## Conflicts of interest

Travel costs for ES to present preliminary results of this study at the 2025 WYMM conference in Hamburg, Germany, were covered by Oxford Nanopore Technologies. RTW received travel expenses from Oxford Nanopore Technologies to attend London Calling 2025. LU has previously received travel expenses from Oxford Nanopore Technologies.

## Funding information

This project was funded by a Helmholtz Principal Investigator Grant awarded to LU. LU was further funded by the University of Zurich. Laboratory and staff costs for routine diagnostics and sample collections were covered by the TUM University hospital and the Institute for Medical Microbiology, Immunology and Hygiene in Munich, Germany and no additional funding was necessary for this study.

## Ethical approval

Ethical approval for the processing of bacterial isolates using nanopore sequencing was given by the TUM ethics committee (2024-522-S-CB). Informed patient consent was waived as samples were taken under routine diagnostics. This research conforms to the principles of the Helsinki declaration.

## Acknowledgements

We thank the laboratory technicians and physicians at the Institute of Medical Microbiology at the Technical University Munich for their support with phenotypic species identification, susceptibility testing, and sample storage and retrieval. This research was made possible by the team responsible for the High-Performance Compute (HPC) platform at Helmholtz Munich and at the New Zealand Institute for Public Health and Forensic Science (PHF Science), with specific thanks to Russell Smithies and Shane Sturrock for HPC.

## Abbreviations

BWA: Burrows-Wheeler Aligner

CFU: colony forming unit

Contig: contiguous sequence

CRE: carbapenem-resistant Enterobacterales

DCJ: Double-Cut-and-Join

ESBL: extended-spectrum beta-lactamase

EUCAST: European Committee on Antimicrobial Susceptibility Testing

GPU: graphics processing unit

GTDB-Tk: Genome Taxonomy Database Toolkit

IPC: Infection Prevention and Control

KPC: *Klebsiella pneumoniae* carbapenemase

MLST: multi-locus sequence typing

NCBI: National Center for Biotechnology Information

PBS: phosphate-buffered saline

SNP: single-nucleotide polymorphism

SRA: Sequence Read Archive

ST: sequence type

TSB: tryptic soy broth

TUM: Technical University of Munich

---

**Supplementary Fig. 1.**
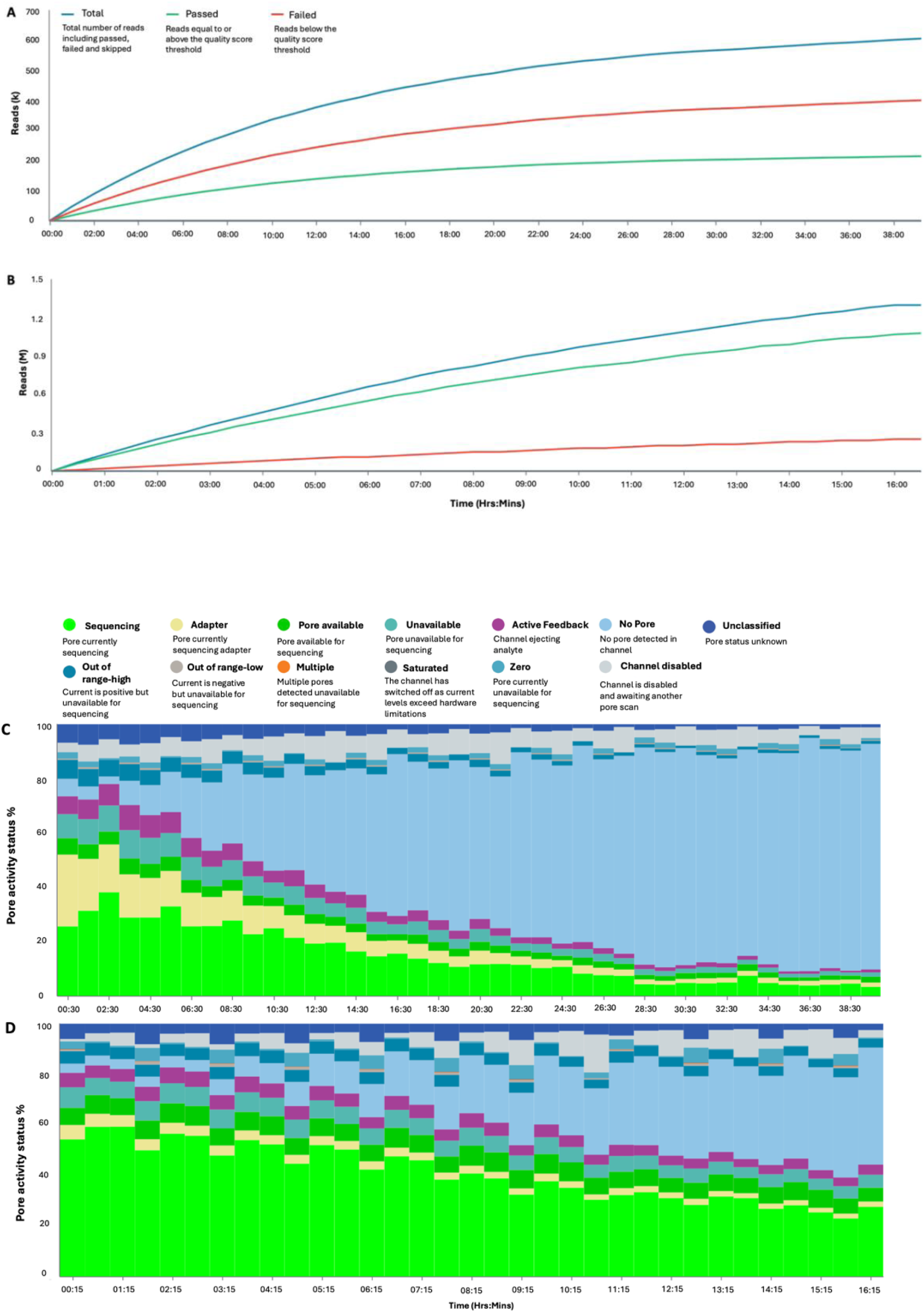
Nanopore sequencing performance comparison without (A, C) and with (B, D) RNase treatment of the same DNA extracts (Table S1; Methods). (A, B) Sequencing read throughput over time, measured by the number of total, passed, and failed reads over sequencing time. Reads are classified as failed if they have a quality score below 8, based on FAST basecalling. Counts area shown in thousands (k, panel A) and millions (M, panel B). (C, D) Nanopore occupancy over sequencing time; actively sequencing pores are colored in bright green, ultimately unavailable pores (for example because of pore clogging) are colored in bright blue and pores sequencing adapters are highlighted in yellow (adapted from the sequencing reports generated by the Oxford Nanopore Technologies’ MinKNOW software v24.06.16).

**Supplementary Fig. 2.**
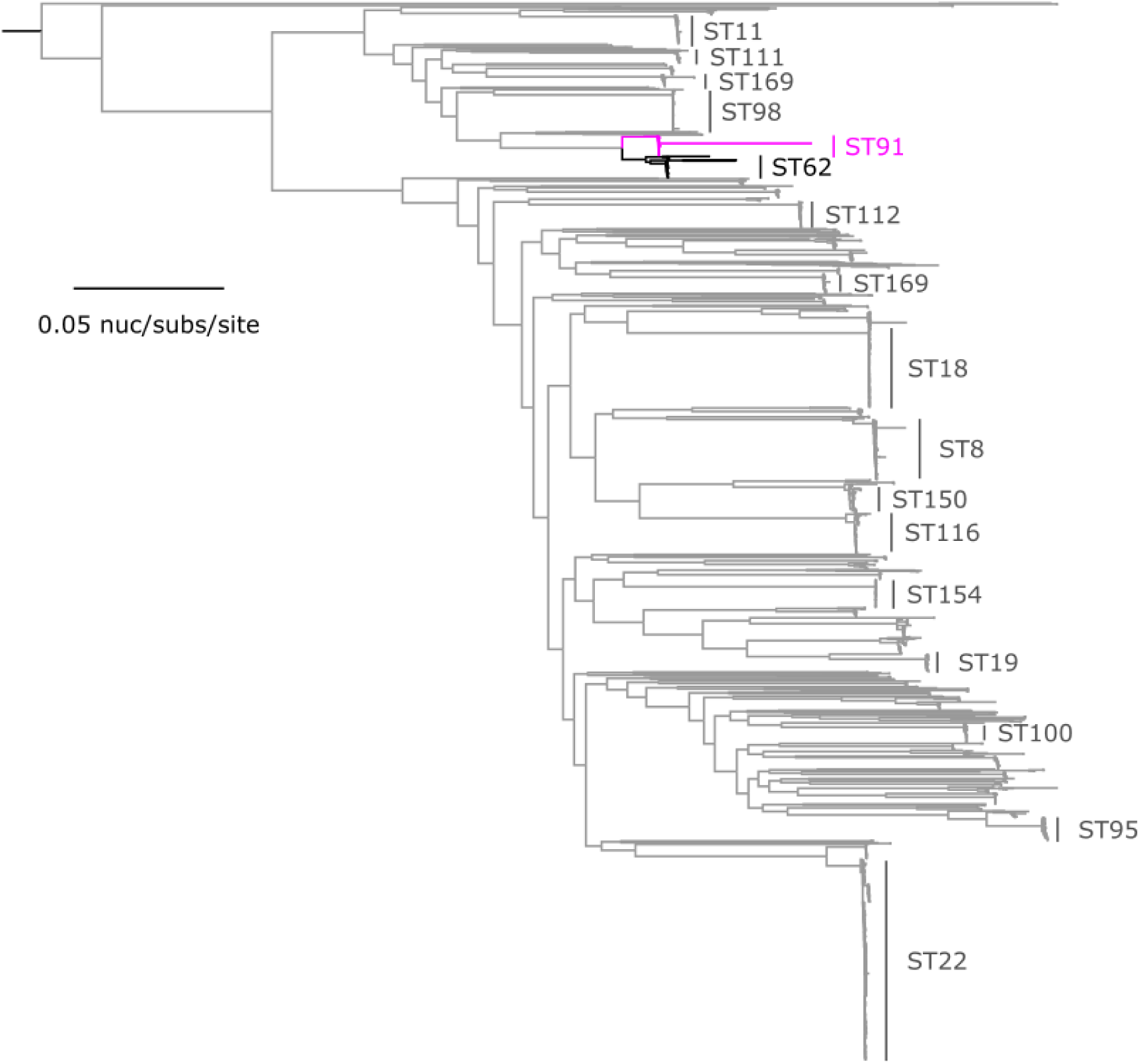
Maximum-likelihood phylogeny of *Citrobacter freundii*. The phylogeny was inferred from 121,511 core-genome single-nucleotide polymorphisms (SNPs) from 2,028 assembled (publicly available) genomes. SNPs were derived from a core-genome alignment of 1,431,649 bp and are called against the 5,093,232 bp complete chromosome of CFTMDU (GenBank: CP151202). A sequence type (ST) comprised of >10 genomes is labelled. The phylogeny was rooted at the midpoint.

**Supplementary Fig. 3.**
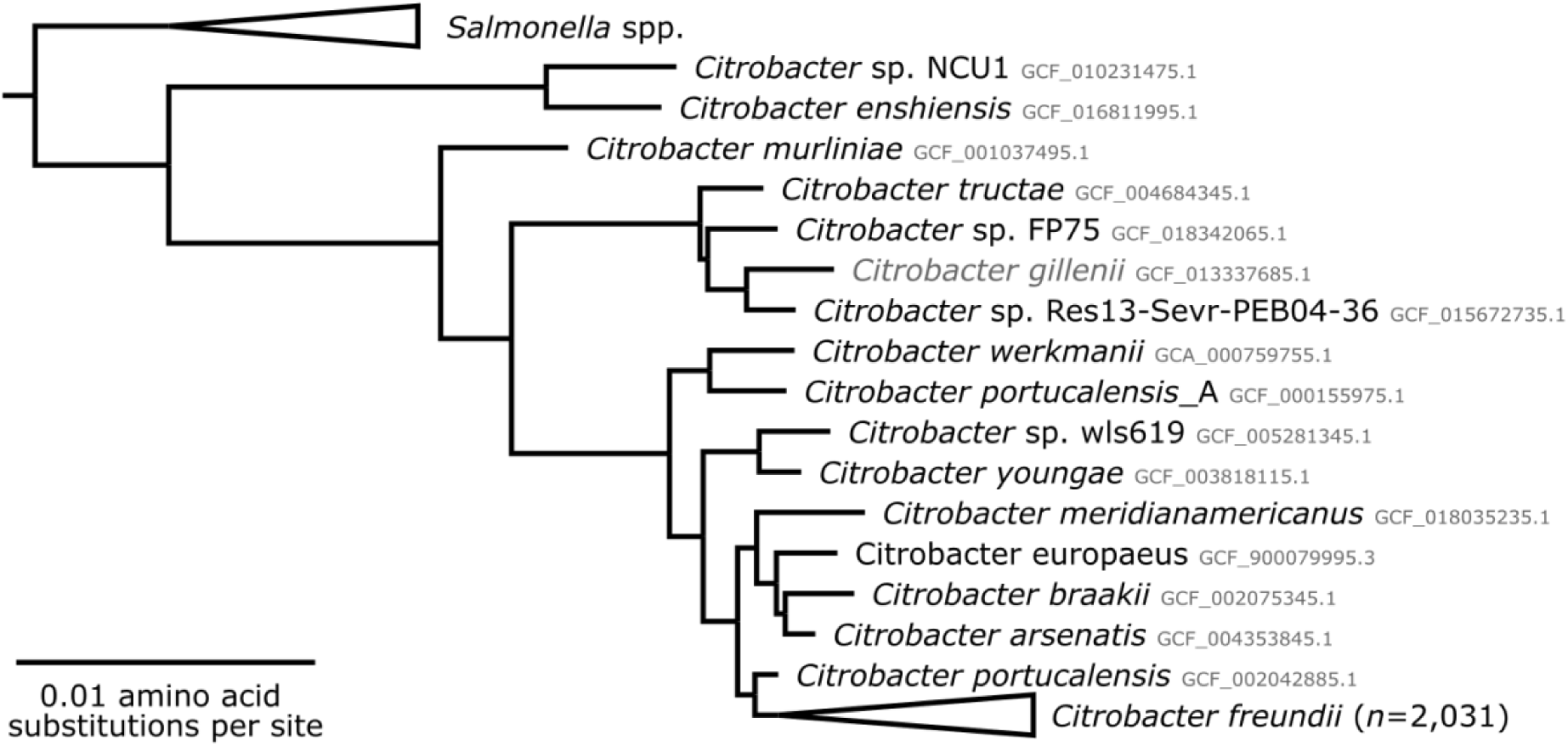
Taxonomic identification of publicly available *C. freundii* complex genomes retrieved from the NCBI nucleotide database and sequence read archive using the GTDB-Tk taxonomic identification tool. The taxonomic tree is based on a maximum-likelihood approximation of protein evolution assuming a gamma-distributed rate heterogeneity. Only genomes that clustered within the *C. freundii* species lineage were included in downstream analyses.

**Supplementary Fig. 4.**
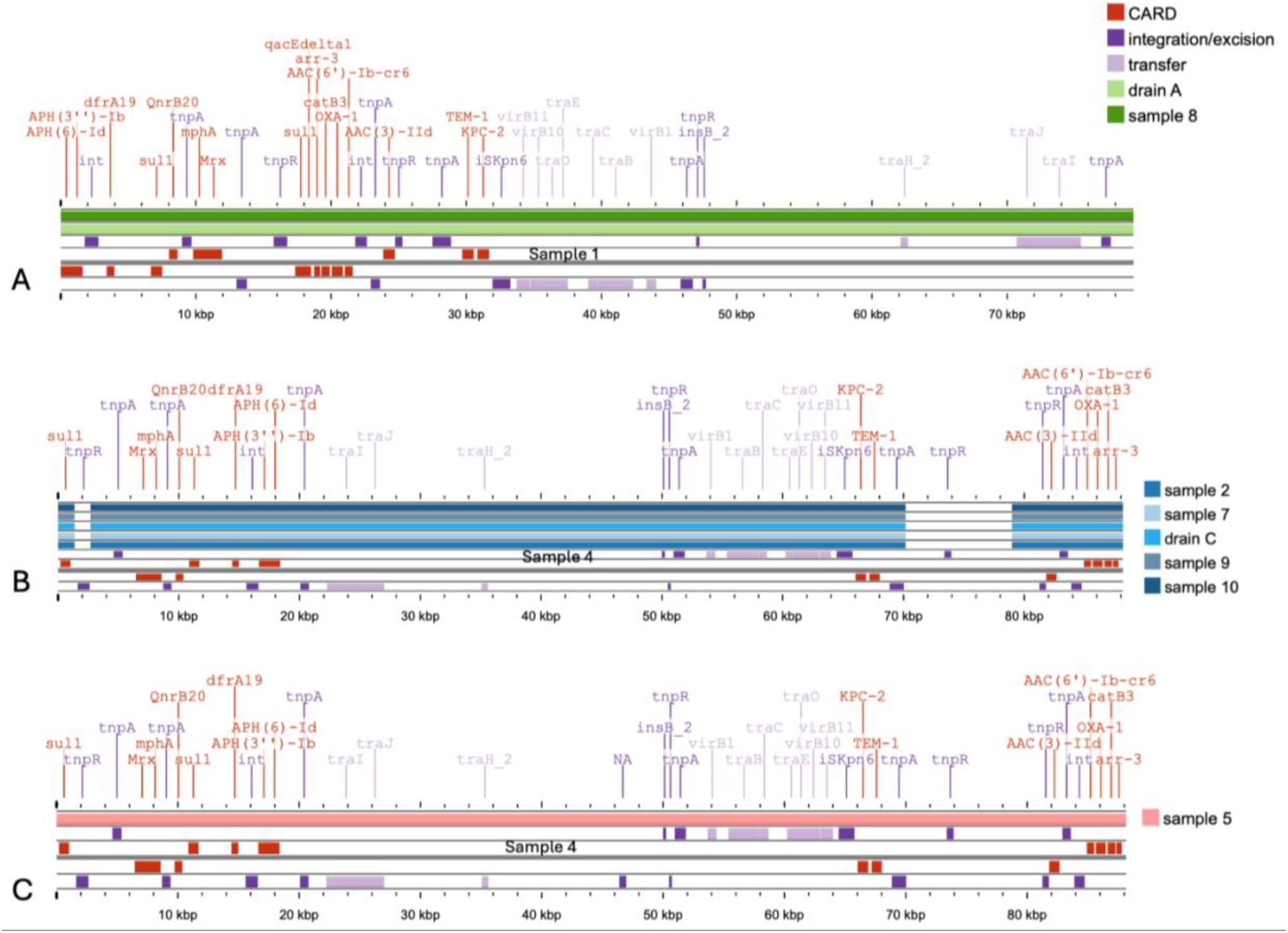
Blastn alignments of IncN plasmids meeting the Mash distance threshold of <0.001 with any other IncN plasmid (A: Sample 1, B: Sample 4, C: Sample 4). The respective chronologically first sample (A, C) or longest sequence (B) are used as reference, respectively, and indicated in grey. Integration and excision annotation based on the mobileOG-db is highlighted in dark purple, and transfer genes based on the mobileOG-db are highlighted in light purple. Created with ProkSee.com

**Supplementary Table 1.**
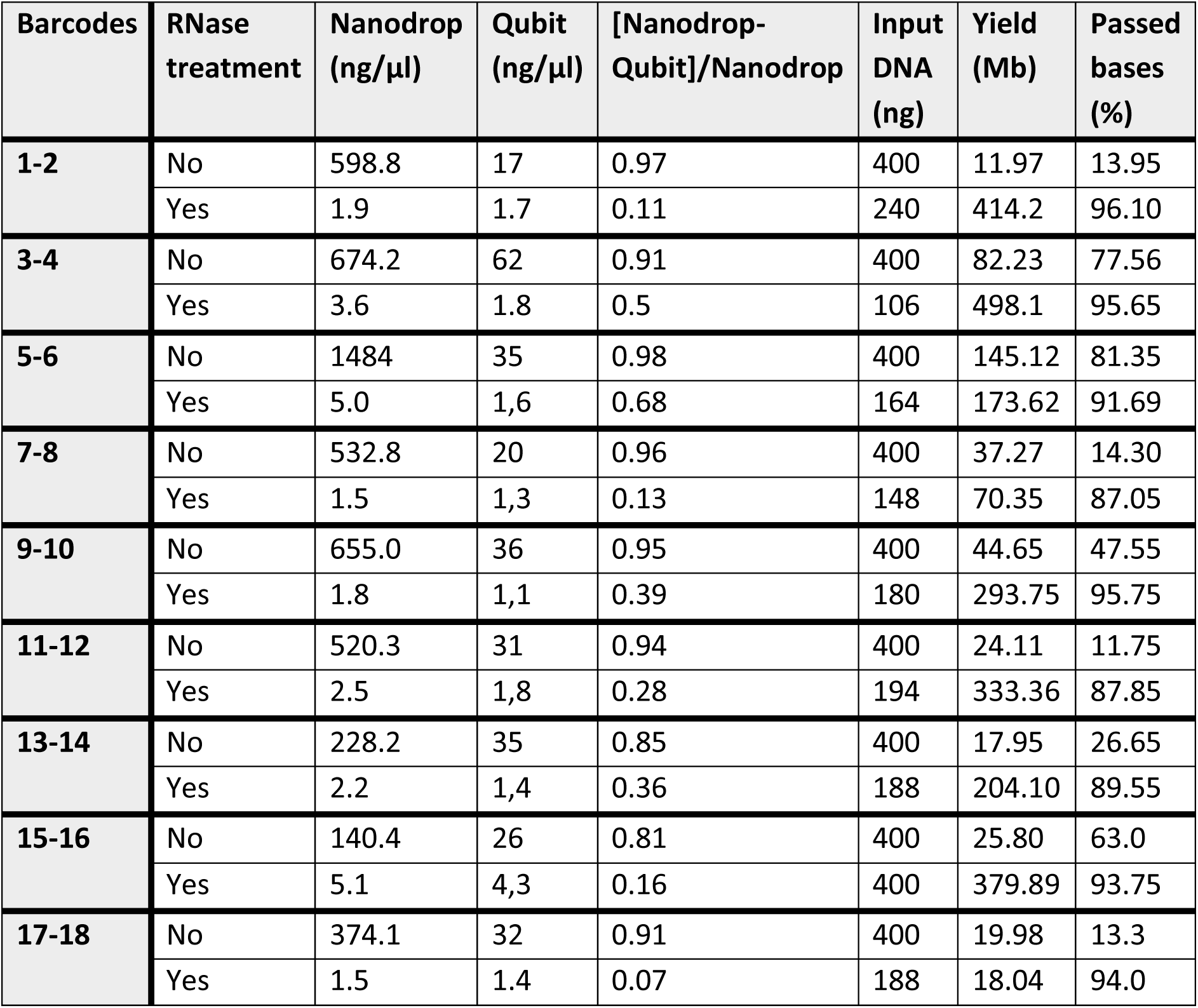
DNA extraction and nanopore sequencing metrics of nine DNA extracts (two barcodes per extracts) that were sequenced without or with post hoc RNase treatment (Methods). Comparison of Nanodrop nucleotide and Qubit DNA concentrations, and their relative difference (measured as [Nanodrop (ng/µl)-Qubit (ng/µl)]/Nanodrop (ng/µl)) prior to upconcentration. Up to 200ng of DNA per barcode were used as input for nanopore library preparation (as recommended by the manufacturer), or as much DNA as available in 10µl of DNA extract after purification and upconcentration. Nanopore sequencing results are presented in total yield (number of Mbases) and percentage of passed bases (after application of a FAST-basecalling quality score filter of 8).

| Barcodes | RNase treatment | Nanodrop (ng/μl) | Qubit (ng/μl) | [Nanodrop-Qubit]/Nanodrop | Input DNA (ng) | Yield (Mb) | Passed bases (%) |
| --- | --- | --- | --- | --- | --- | --- | --- |
| **1-2** | No | 598.8 | 17 | 0.97 | 400 | 11.97 | 13.95 |
|  | Yes | 1.9 | 1.7 | 0.11 | 240 | 414.2 | 96.10 |
| **3-4** | No | 674.2 | 62 | 0.91 | 400 | 82.23 | 77.56 |
|  | Yes | 3.6 | 1.8 | 0.5 | 106 | 498.1 | 95.65 |
| **5-6** | No | 1484 | 35 | 0.98 | 400 | 145.12 | 81.35 |
|  | Yes | 5.0 | 1,6 | 0.68 | 164 | 173.62 | 91.69 |
| **7-8** | No | 532.8 | 20 | 0.96 | 400 | 37.27 | 14.30 |
|  | Yes | 1.5 | 1,3 | 0.13 | 148 | 70.35 | 87.05 |
| **9-10** | No | 655.0 | 36 | 0.95 | 400 | 44.65 | 47.55 |
|  | Yes | 1.8 | 1,1 | 0.39 | 180 | 293.75 | 95.75 |
| **11-12** | No | 520.3 | 31 | 0.94 | 400 | 24.11 | 11.75 |
|  | Yes | 2.5 | 1,8 | 0.28 | 194 | 333.36 | 87.85 |
| **13-14** | No | 228.2 | 35 | 0.85 | 400 | 17.95 | 26.65 |
|  | Yes | 2.2 | 1,4 | 0.36 | 188 | 204.10 | 89.55 |
| **15-16** | No | 140.4 | 26 | 0.81 | 400 | 25.80 | 63.0 |
|  | Yes | 5.1 | 4,3 | 0.16 | 400 | 379.89 | 93.75 |
| **17-18** | No | 374.1 | 32 | 0.91 | 400 | 19.98 | 13.3 |
|  | Yes | 1.5 | 1.4 | 0.07 | 188 | 18.04 | 94.0 |

**Supplementary Table 2.** Sequencing summary of the filtered and basecalled nanopore sequencing data across all isolates, including total number of reads, read-level N50, total number of bases, as well as mean base-level quality score, and percentage of bases with a quality score above 20.

| Sample | # reads | read N50 | # bases | mean Q-score | Q20 (%) |
| --- | --- | --- | --- | --- | --- |
| **1** | 130,785 | 5,801 | 456,322,626 | 22.16 | 94.85 |
| **2** | 166,338 | 5,228 | 496,481,848 | 21.43 | 93.91 |
| **3** | 99,683 | 6,402 | 358,492,576 | 20.64 | 92.26 |
| **4** | 189,570 | 6,897 | 690,118,598 | 20.98 | 93.03 |
| **5** | 500,513 | 6,328 | 1,769,558,787 | 21.4 | 93.74 |
| **6** | 128,310 | 9,480 | 593,636,606 | 21.0 | 93.17 |
| **7** | 130,553 | 4,272 | 321,674,712 | 20.42 | 92.92 |
| **8** | 265,115 | 10,340 | 1,406,718,773 | 21.84 | 94.36 |
| **9** | 269,725 | 6,665 | 944,512,723 | 21.21 | 93.72 |
| **10** | 180,221 | 8,133 | 713,718,257 | 21.58 | 93.98 |
| **Drain A** | 322,607 | 8,139 | 1,353,038,408 | 21.83 | 94.26 |
| **Drain B** | 113,529 | 5,573 | 382,614,256 | 21.40 | 94.00 |
| **Drain C** | 182,033 | 8,897 | 827,156,867 | 21.08 | 93.22 |

**Supplementary Table 3.** Assembly metrics overview of each isolates’ bacterial chromosome and (if present) IncN plasmid, including contig length, median coverage, and contig circularity. No IncN plasmid was identified in the annotated assemblies from sample 6.

| Sample | Chromosome |  |  | IncN plasmid |  |  |
| --- | --- | --- | --- | --- | --- | --- |
|  | Contig length (b) | Median coverage | Circular | Contig length (b) | Median coverage | Circular |
| **1** | 4,904,886 | 85 | ✓ | 79,348 | 162 | ✓ |
| **2** | 5,248,615 | 90 | ✓ | 78,021 | 156 | ✓ |
| **3** | 5,134,109 | 59 | ✓ | 72,036 | 132 | ✓ |
| **4** | 5,247,584 | 58 | ✓ | 88,158 | 41 | ✓ |
| **5** | 5,131,833 | 314 | ✓ | 88,157 | 273 | ✓ |
| **6** | 5,303,591 | 46 | X | NA | NA | NA |
| **7** | 5,248,812 | 109 | ✓ | 78,051 | 179 | ✓ |
| **8** | 4,860,193 | 282 | ✓ | 79,348 | 386 | ✓ |
| **9** | 5,248,611 | 107 | ✓ | 77,380 | 214 | ✓ |
| **10** | 5,348,650 | 70 | ✓ | 77,391 | 85 | ✓ |
| **Drain A** | 4,794,503 | 275 | ✓ | 79,349 | 263 | ✓ |
| **Drain B** | 5,020,617 | 57 | ✓ | 81,368 | 64 | ✓ |
| **Drain C** | 5,249,966 | 152 | ✓ | 78,032 | 253 | ✓ |

**Supplementary Table 4.**
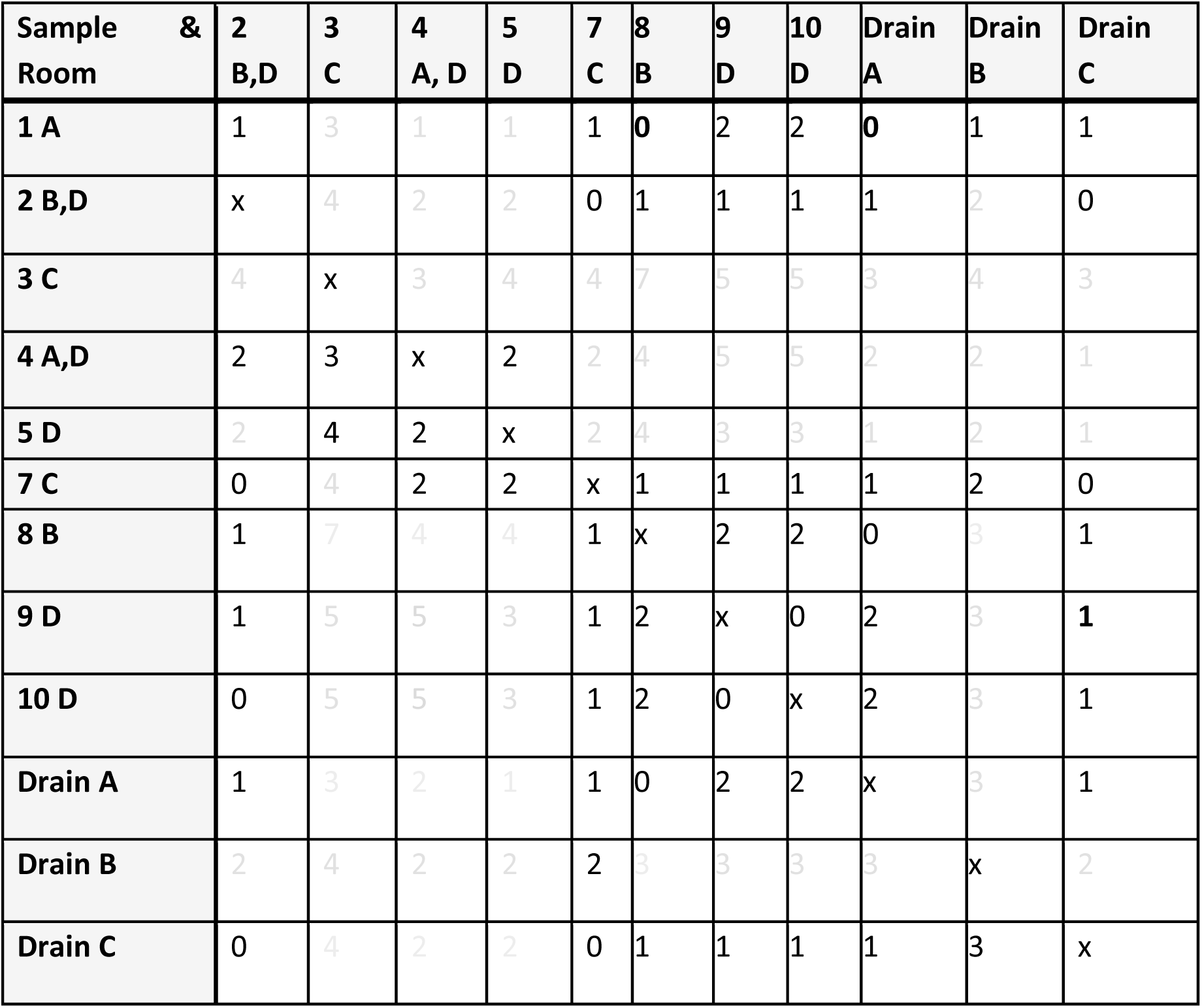
Contingency table of DCJ-indel distances between all IncN plasmids. Isolates are described by their sample number and associated hospital rooms. The plasmid pairs that did not meet the Mash threshold of <0.001 are marked in grey (Methods).

| Sample &<br>Room | 2<br>B,D | 3<br>C | 4<br>A, D | 5<br>D | 7<br>C | 8<br>B | 9<br>D | 10<br>D | Drain<br>A | Drain<br>B | Drain<br>C |
| --- | --- | --- | --- | --- | --- | --- | --- | --- | --- | --- | --- |
| 1 A | 1 | 3 | 1 | 1 | 1 | 0 | 2 | 2 | 0 | 1 | 1 |
| 2 B,D | x | 4 | 2 | 2 | 0 | 1 | 1 | 1 | 1 | 2 | 0 |
| 3 C | 4 | x | 3 | 4 | 4 | 7 | 5 | 5 | 3 | 4 | 3 |
| 4 A,D | 2 | 3 | x | 2 | 2 | 4 | 5 | 5 | 2 | 2 | 1 |
| 5 D | 2 | 4 | 2 | x | 2 | 4 | 3 | 3 | 1 | 2 | 1 |
| 7 C | 0 | 4 | 2 | 2 | x | 1 | 1 | 1 | 1 | 2 | 0 |
| 8 B | 1 | 7 | 4 | 4 | 1 | x | 2 | 2 | 0 | 3 | 1 |
| 9 D | 1 | 5 | 5 | 3 | 1 | 2 | x | 0 | 2 | 3 | 1 |
| 10 D | 0 | 5 | 5 | 3 | 1 | 2 | 0 | x | 2 | 3 | 1 |
| Drain A | 1 | 3 | 2 | 1 | 1 | 0 | 2 | 2 | x | 3 | 1 |
| Drain B | 2 | 4 | 2 | 2 | 2 | 3 | 3 | 3 | 3 | x | 2 |
| Drain C | 0 | 4 | 2 | 2 | 0 | 1 | 1 | 1 | 1 | 3 | x |

## Notes

### Competing Interest Statement

The authors have declared no competing interest.

